# Stress Evaluation of Various Breeding Environments in Mouse

**DOI:** 10.1101/2021.08.27.457934

**Authors:** Gwang-Hoon Lee, Hye-Yoon Choi, Woori Jo, Kil-Soo Kim

## Abstract

Laboratory animals are raised in a fixed space during the study period and are environmentally bound. Laboratory animal may be under stress on the constrained environment, which changes physiological indicators, affecting the reproducibility and accuracy of animal study. Therefore, reducing animal stress by providing proper breeding environment and environmental enrichment can be the basis for animal study. In this study, the stress level was assessed according to the mouse breeding environment. According to the results of the experiment, it was determined that the individual ventilation cage had less cortisol concentration in serum and body weight increased in the individual ventilation cage than individual isolated cage, when providing environmental enrichment rather than group breeding or not providing environmental enrichment. The results will provide appropriate guidelines for laboratory animal welfare.

## Introduction

With the recent increase in interest in animal ethics, there has been a growing interest in improvement of housing environment and animal welfare in the international laboratory animal research community (Hagelin, Hau et al. 2003). Although related laws and systems have been continuously strengthened in the field of the laboratory animal studies, issues about ethical relations humans and animal in bioscience research laboratories continue. The Ministry of Agriculture, Food and Rural Affairs (MAFRA, Republic of Korea) announced a five-year comprehensive plan for animal welfare in order to raise public awareness and sympathy for animal protection and welfare, enhance animal laboratory animal ethics by reinforcing the function of IACUC. Recently, furthermore, discussions on the laboratory animal welfare are underway at the National Assembly debate.

In addition, especially unlike domestic companion animals, laboratory animals are limited to a fixed space for the entire experimental period, so laboratory animals are more exposed to stress from housing environment than companion animals. This constrained environmental condition can cause stress on animals, which affects the physiological indicators of animals and can change the results of the experiments. Therefore, in order to obtain reliable experimental results in the laboratory animal research, animal welfare is important to alleviate the stress of laboratory animals because it affects the experimental results (Davies, Greenhough et al. 2018). Also, policies and practice with respect to laboratory animal housing, husbandry, and quality care can enhance animal welfare (Kappel, Hawkins et al. 2017)

Laboratory animals are used in a variety of research fields, and stress induced during experiments can change background data. Stress is defined as the body’s non-specific response to external stimuli, environmental demands, or stimuli beyond the body’s ability to cope with them (Fink 2016). Vladimir K. et al. reported that stress change limbic-hypothalamo-pituitary-adrenal (LHPA) neuroendocrine axis and De Kloet ER et al reported stress induced structural and functional change in limbic brain and B Olivier demonstrated that stress increased body temperature of laboratory animal (GP and Gold 1992, Olivier, Zethof et al. 2003, Patchev and Patchev 2006).

Most studies on stress are the experiments with artificially applied stress including repeated social defeat stress (RSDS), electric shock, wire netting, repeated stress, etc (Rybkin, Zhou et al. 1997, Harris, Zhou et al. 1998, Kim, Jung et al. 1998, Gong, Miao et al. 2015, Takahashi, Chung et al. 2017). In addition, previous studies were performed with efficacy studies of new drug development using artificially inducing stress model (Kim, Jung et al. 1998, Takahashi, Chung et al. 2017), hormonal changes caused by different levels of stress exposure (Alfarez, Wiegert et al. 2002). However, studies on the stress level exposed to various housing environments for improvement of laboratory animal welfare are lacking. Furthermore, it is necessary to evaluate environmental effects on stress of laboratory animals depending on the type of cage, and/or social isolation stress.

Stress-response hormones include cortisol and corticosterone; cortisol is widely used in studies with beagles of large laboratory animals. (Coppola, Grandin et al. 2006, Uccheddu, Mariti et al. 2018), but in rodents, corticosterone is more important glucocorticoid responding to stress exposure (Gong, Miao et al. 2015).

Therefore, this study aimed to evaluate changes in serum corticosterone concentration in serum and body weight as stress indicators according to the presence or absence of environmental enrichment, different types of cages (IVC, individually ventilated cages and ISO, individually isolated cages), and social isolation stress (single breeding and group breeding). These results would be used to enhance environmental conditions for laboratory animals and improve animal ethics by reducing stress on laboratory animals during entire experimental period.

## Materials and Methods

### Animals and Husbandry

A four-week-old CrljOri:CD1(ICR) mouse was purchased from Orient Bio (Seongnam, KyungKi, Korea). The animal experiments were reviewed and approved by the Institutional Animal Care and Use Committee at the Daegu-Gyeongbuk Medical Innovation Foundation (Approved IACUC Number: DGMIF-20032407, Approved date: 23 march 2020). Animals were fed an autoclaved pellet diet (SAFE+40RMM; SAFE Diets, Augy, France) and provided drinking water *ad libitum*. Animals were hosed in environmental conditions with a temperature of 22 ± 1°C, a humidity of 50 ± 10%, illumination at 150-300 Lux, and a ventilation cycle of breeding room was 10-20 times/h. All animals were monitored every day and there is no mortality injury and any clinical signs.

### Experimental Design

The Experimental design is shown in Figure 1A. The mice were divided into six groups (nine mice/group) as follows: (A) IVC/Single (B) IVC/Single + E.E (C) IVC/Group (D) IVC/Group + E.E (E) ISO/Single (F) ISO/Group. IVC (Cat No: GM500) and ISO (Cat No. ISO cage-N). Both cage systems (Tecniplast, Buguggiate, Varese, Italy) have an air circulation system in each individual cage, but the main difference is whether external air is allowed to enter or not. In IVC cage, the external air can enter through the HEPA filter (Size: 141 × 170 mm, Efficiency: 0.3micrometer of particle at 99.5%) on the IVC lid, but in the ISO cage (Size: 73 × 73 × 24 mm, Efficiency: 0.3micrometer of particle at 99.97%) outside air is completely blocked.

**Figure 1.**
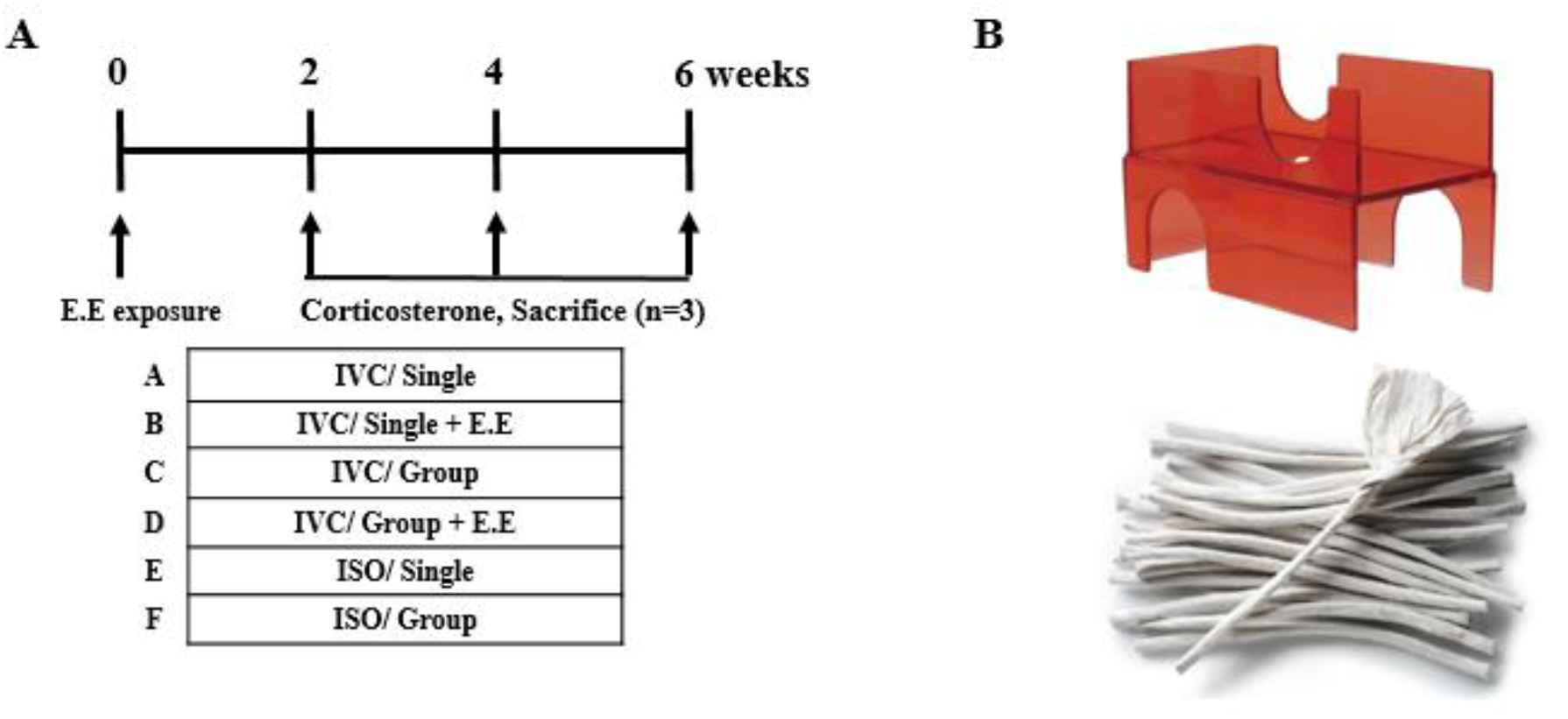
Experimental design (A) and the representative photographs of environmental enrichment (B, harbor mouse retreat, top; diamond twist, bottom)

Two environmental enrichment (E.E) materials were used, the first being harbor mouse retreat (Figure 1B top, Cat No. K3583, Bio-serv, Frenchtown, NJ, USA) and the second being a diamond twist (Figure 1B bottom, Envigo, Madison, WI, USA), which were provided to groups B and D until the end of the experiment. Harbor mouse retreat and diamond twist were selected for E.E materials as standard environmental enrichment required and additional enrichment, respectively with reference to IACUC policy of University of California, Irvine.

### Preparation of blood serum and corticosterone assay

In the present study, the blood collections were performed once every 2 weeks, 3 mice in each group. Mice were anesthetized using isoflurane at 17:00-18:00 pm and blood collection was rapidly performed to minimize the stress caused by the anesthesia process and decapitated. Blood was collected from abdominal vein (about 600 ul) and in serum separate tubes (SST tube, Becton, Dickinson and Company, Franklin Lakes, NJ, USA), centrifuged (3000 rpm, 10 min, 4°C) to separate serum.

Triplicate serum corticosterone assay was conducted using ELISA kit (Cat No. K014, Arbor assays, Ann Arbor, Michigan, USA) and optical density (OD) read by synergy H4 microplate reader (BioTek Instruments, Inc. Winooski, VT, USA)

### Statistical Analysis

Statistical significance was determined using GraphPad Prism 8 (GraphPad Software Inc., San Diego, CA, USA). All data are presented as mean ± standard error of the mean (SEM) and for comparisons among each groups, two-way ANOVA with Tukey’s multiple comparisons test and Student’s t-test were used (Figure 2).

**Figure 2.**
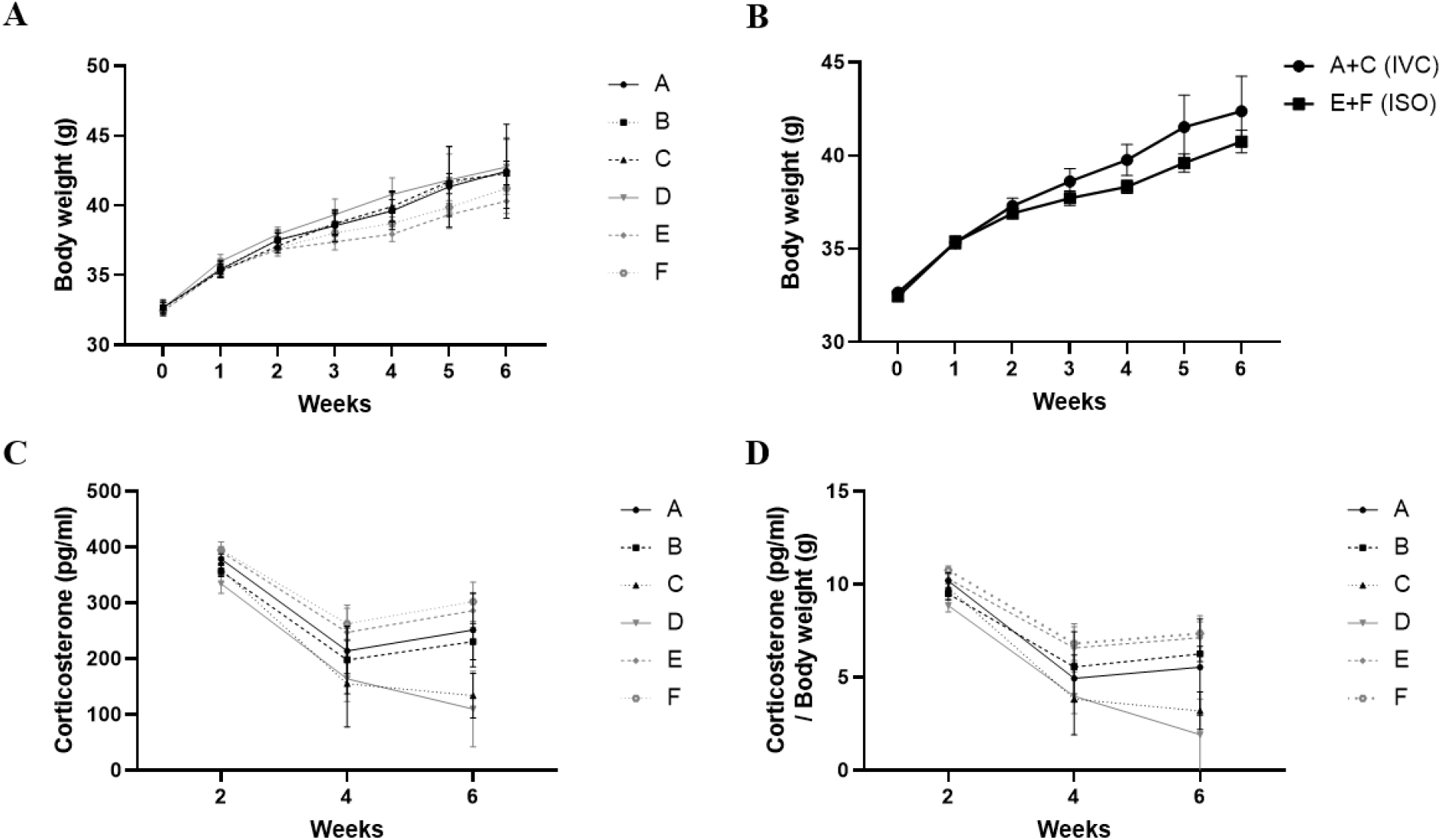
Body weight(g) in each group (A) and in group A+C and E+F (B) and concentrations of serum corticosterone in mice at different times of week.

## Results

### Body weight

Nine mice per group were sacrificed three every two weeks. After a week of acclimatization, body weight was measured once a week until the euthanasia. During the experiment period, group D had the highest average body weight, followed by group A, C, B, F, E. There was a significant difference between groups C and F, groups A and E, and groups B and D by the Students T-test.

### Concentrations of serum corticosterone

From 2 to 4 weeks, concentrations in serum corticosterone in all groups decreased, and from 4 to 6 weeks, group C and D, which had been group reared, decreased, and the remaining groups increased slightly again. Serum corticosterone concentration shows the order of F, E, A, B, C, D at week 6, and the relative corticosterone concentration (corticosterone / body weight) has an overall similar tendency to the serum corticosterone level, but there is a difference that group A value is lower than B value. In groups A and B, corticosterone level showed statistical significance with the Students T-test.

## Discussions

With the revision of the Animal Protection Act in 2007 in Republic of Korea, the “principle of 3R” emerged as a principle in laboratory animal research and Institutional Animal Care and Use Committee widely began to be launched. Also, recently, response-ability as the idea of a fourth R is emerging as important beyond the 3R which are replacement as much as possible, reduce the number of animals, and refinement to animal pain, stress in the course of experiments(Davies, Greenhough et al. 2018). Therefore, it is necessary to establish a method that induces less stress while improving the laboratory animal welfare, and related studies are needed accordingly.

Stress affects mobilization of physical and psychological tension for coping with difficult situation, adversity, and restoring homeostasis; in addition, when stress is induced, it affects the reproducibility and reliability of laboratory animal research (Armario, Garcia-Marquez et al. 1987, Bligh, DeStefano et al. 1990, Armario, Martí et al. 1993, Armario, Gavaldà et al. 1995).

In order to improve the reproducibility and reliability of experiments, the DGMIF laboratory animal center evaluates the level of stress induced by the environment of laboratory animals and provides high-quality animal experiments by realizing animal welfare.

In this study, we provided the 6 different environmental conditions to the mice in groups raised at the DGMIF laboratory animal center to evaluate the level of the stress. Two environmental enrichment materials provided, a house for mouse environmental environment where the mouse can hide or run up and a Diamond twist that can be gnawed with the mouse teeth to form a nest.

Previous studies showed that the relationship between stress and behavioral changes(Fernandez, Collazo et al. 2004, Mineur, Belzung et al. 2006, Uccheddu, Mariti et al. 2018, Liu, Wang et al. 2019) For example, the unpredictable chronic mild stress (UCMS) induced by cage tilting, damp saw dust, predator sounds, placement in an empty cage, placement in an empty cage with water on the bottom, inversion of the light/dark cycle, lights on for a short period of time during the dark phase, or switching cages induces increased depressive-like behaviors (Mineur, Belzung et al. 2006).

In addition, there are some studies about the effectiveness of E.E. (Benaroya-Milshtein, Hollander et al., Fernandez, Collazo et al. 2004, Rehnberg, Robert et al. 2015, Uccheddu, Mariti et al. 2018), which prevents oxidative injury and restores cholinergic neurotransmission in cognitively impaired aged rat(Fernandez, Collazo et al. 2004), also alleviates the behavioral changes in mouse model of posttraumatic stress disorder(Golub, Kaltwasser et al. 2011). Also, the studies about the relationship between housing condition such as single housing or grouped housing and the behavioral phenotype were conducted (Kulesskaya, Rauvala et al. 2011), both isolation and environmental enrichment have fundamental effects on mouse behavior and should be considered in the course of experimental design with stress-related animal models and animal welfare assessment.

In this study, body weight can be a good indicator to evaluate stress level, and it is generally known that the amount of feed intake and body weight decreases, the level of stress can be an indicator of body weight (Rybkin, Zhou et al. 1997, Harris, Zhou et al. 1998, Takahashi, Chung et al. 2017). This study reports that animal exposed to stress might decrease body weight gain rate. Based on the results of body weight in groups, group D (IVC cage/Group + E.E), which has increased the most, is less stressful, and group E (ISO cage/single), which has gained the least weight, represents the status under more stressful environment (Table1). Corticosterone in the blood is widely known to be an increasing factor when the mouse is exposed to stress. (Gong, Miao et al. 2015). Therefore, based on the results of serum corticosterone concentration, group D (IVC cage/Group+E.E) with the lowest concentration was the least stressed, and group F (ISO cage/Group) with the highest concentration was the most stressed. Overall, the concentration of corticosterone in serum was the highest at the second week, which was determined to be the highest at 2-weeks and then decreased. The results of the relative serum corticosterone (corticosterone / body weight) was generally similar to the result of serum corticosterone, only changing the order of group A (IVC/Single) and group B (IVC/Single + E.E).

**Table 1.**
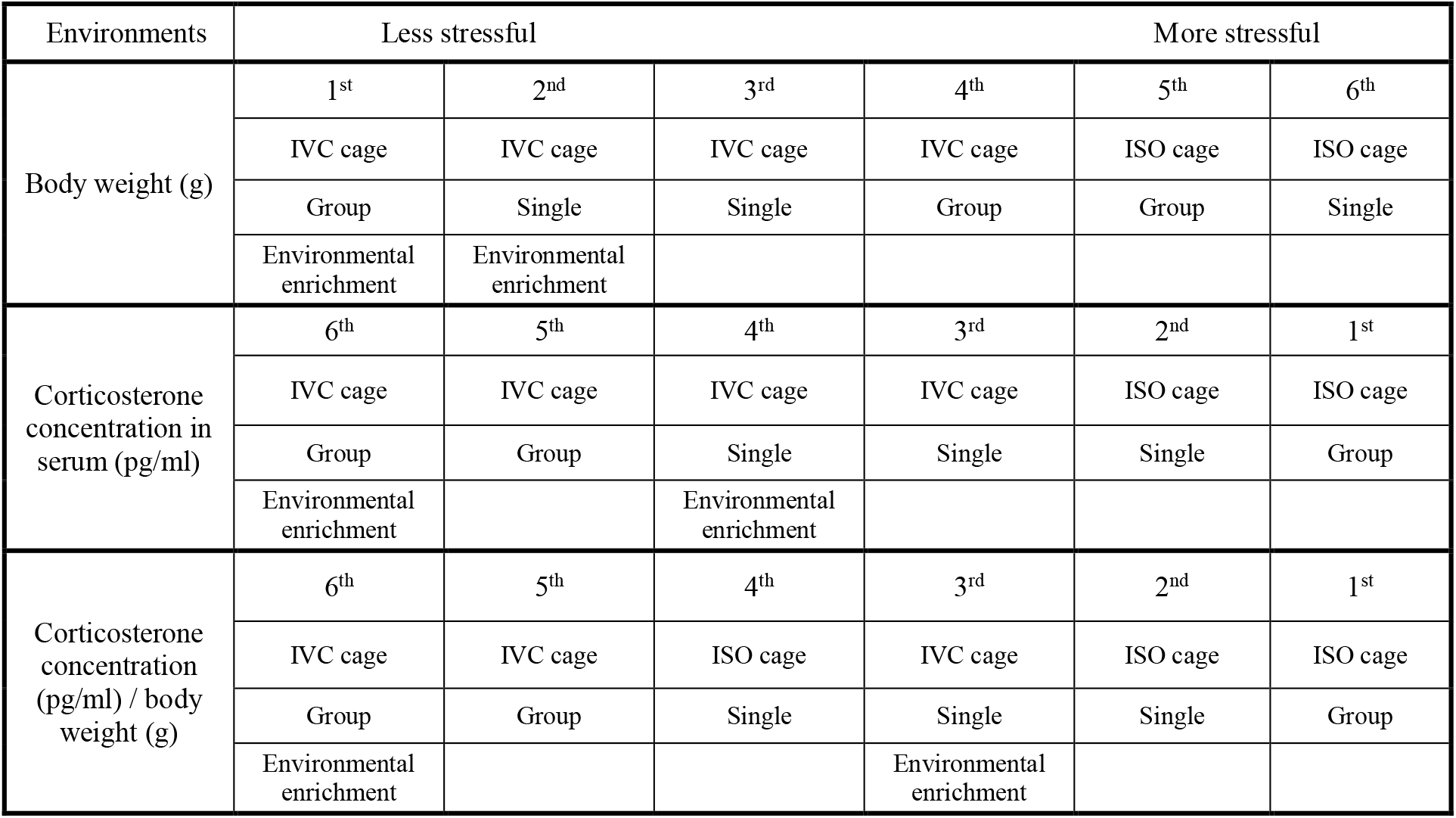
Overall analysis for body weight and corticosterone concentration in serum and corticosterone concentration / body weight

| Environments | Less stressful |  |  | More stressful |  |  |
| --- | --- | --- | --- | --- | --- | --- |
| Body weight (g) | 1<sup>st</sup> | 2<sup>nd</sup> | 3<sup>rd</sup> | 4<sup>th</sup> | 5<sup>th</sup> | 6<sup>th</sup> |
|  | IVC cage | IVC cage | IVC cage | IVC cage | ISO cage | ISO cage |
|  | Group | Single | Single | Group | Group | Single |
|  | Environmental enrichment | Environmental enrichment |  |  |  |  |
| Corticosterone concentration in serum (pg/ml) | 6<sup>th</sup> | 5<sup>th</sup> | 4<sup>th</sup> | 3<sup>rd</sup> | 2<sup>nd</sup> | 1<sup>st</sup> |
|  | IVC cage | IVC cage | IVC cage | IVC cage | ISO cage | ISO cage |
|  | Group | Group | Single | Single | Single | Group |
|  | Environmental enrichment |  | Environmental enrichment |  |  |  |
| Corticosterone concentration (pg/ml) / body weight (g) | 6<sup>th</sup> | 5<sup>th</sup> | 4<sup>th</sup> | 3<sup>rd</sup> | 2<sup>nd</sup> | 1<sup>st</sup> |
|  | IVC cage | IVC cage | ISO cage | IVC cage | ISO cage | ISO cage |
|  | Group | Group | Single | Single | Single | Group |
|  | Environmental enrichment |  |  | Environmental enrichment |  |  |

Based on these results, Group D (IVC/Group + E.E) was the most weighted group, and the corticosterone concentration was the lowest on average, so it was determined that stress was the least. Also, the groups with E.E show less stressful environment than the groups without E.E. However, there was no difference in body weight or corticosterone concentration between group breeding and single breeding. However, it was difficult to judge the results of the relationship between group housing and single housing condition, and in our experiment, it was not enough to consider a sufficient number of individuals. In addition, there is a divided opinion as things due to social ranking can also be a hindrance to judging the results. Benaroya-Milshtein, Hollander et al. reported in the mouse that the E.E reduces anxiety and weakens stress reactions, while Chapillo, Pet al. reported that the Environmental energy in the mouse reduced anxiety (Benaroya-Milshtein, Hollander et al., Chapillon, Manneche et al. 1999). Although the mouse is a group animal, it is judged that there is a difference in the animal because it can be exposed to stress due to the hierarchy that is determined by the group. Both group and single breeding are known to have pros and cons, and the results of various experiments showed conflicting results. (Kappel, Hawkins et al. 2017). Liu, Wang et al. reported that single in mice reduced the concentration of corticosterone and kamakura, Kovalainen et al. reported that the group had higher levels of corticosterone than single in mice, indicating that the group had higher stress (Kamakura, Kovalainen et al. 2016, Liu, Wang et al. 2019). On the other hand, Norman, K. J., et al. reported that single significantly reduced the memory of the mouse, and according to a study by Kamle, A et al., increased the level of LTP injury in the single breeding group hippocampus in the C57BL/6J mouse and had higher blood Corticosterone concentration (Kamal, Ramakers et al. 2014, Norman, Seiden et al. 2015).

In addition, in this study, the average weight of ISO cage is lower than IVC cage and the blood corticosterone concentration is higher on average, so it is judged that the stress is higher because ISO cage has limited air circulation.

Furthermore, stress effects may be different for each mouse strain, so more studies on stress in diverse strains and species using animal models with disease are needed (Olivier, Zethof et al. 2003). Also, with the growing interest in animal welfare, stress assessment could be expanded in abandoned animals or industrial animals (Rehnberg, Robert et al. 2015).

DGMIF laboratory animal center was certified by the Ministry of Food and Drug Safety in 2016 and was award full accreditation by the Association for Assessment and Accreditation of Laboratory Animal Care International (AAALAC-i) in 2020. Full accreditation of AAALAC-i is a certification that is recognized as a non-clinical study institution that encourages humane treatment of animals in the field of science and that it maintains a high level for laboratory animal care and use (Gettayacamin and Retnam 2017). In addition to this full accreditation of AAALAC-i, the DGMIF Laboratory animal center is devising and continuing to develop ways to promote welfare for Laboratory animals.

These results suggest that IVC cages are preferred than ISO cages, and providing an E.E would be better to relieve stress. However, proper animal composition is needed because stress can be more active depending on animals raised in the same cage. This study will be helpful to provide appropriate guidelines for the management and operation of laboratory animal breeding and the welfare of laboratory animals.

## Abbreviation

E.E: Environmental Enrichment
IVCs: Individually Ventilated Cages
ISO: Individually Isolated Cages

